# ATXN10 is required for embryonic heart development and maintenance of epithelial cell phenotypes in the adult kidney and pancreas

**DOI:** 10.1101/2021.04.29.441883

**Authors:** Melissa R. Bentley-Ford, Reagan S. Andersen, Mandy J. Croyle, Courtney J. Haycraft, Kelsey R. Clearman, Jeremy B. Foote, Jeremy F. Reiter, Bradley K. Yoder

## Abstract

*Atxn10* is a gene known for its role in cytokinesis during the cell cycle and is associated with Spinocerebellar Ataxia (SCA10), a slowly progressing cerebellar syndrome caused by an intragenic pentanucleotide repeat expansion. *Atxn10* is also implicated in the ciliopathy syndromes Nephronophthisis (NPHP) and Joubert Syndrome (JBTS), which are caused by disruption of cilia function leading to nephron loss, impaired renal function, and cerebellar hypoplasia. How *Atxn10* disruption contributes to these disorders remains unknown. Here we generated *Atxn10* congenital and conditional mutant mouse models. Our data indicate that while ATXN10 protein can be detected around the base of the cilium as well as in the cytosol, its loss does not cause overt changes in cilia formation or morphology. Congenital loss of *Atxn10* results in embryonic lethality around E10.5 associated with pericardial effusion and loss of trabeculation. Similarly, tissue specific loss of ATXN10 in the developing endothelium (Tie2-Cre) and myocardium (cTnT-Cre) also results in embryonic lethality with severe cardiac malformations occurring in the latter. Using an inducible Cagg-CreER to disrupt Atxn10 systemically, we show that ATXN10 is also required for survival in adult mice. Loss of ATXN10 results in severe pancreatic and renal abnormalities leading to lethality within a few weeks post ATXN10 deletion in adult mice. Evaluation of these phenotypes further identified rapid epithelial to mesenchymal transition (EMT) in these tissues. In the pancreas, the phenotype includes signs of both acinar to ductal metaplasia and EMT with aberrant cilia formation and severe defects in glucose homeostasis related to pancreatic insufficiency or defects in feeding or nutrient intake. Collectively this study identifies ATXN10 as an essential protein for survival.

## Introduction

*Ataxin10* (ATXN10) is most commonly associated with spinocerebellar ataxia type 10 (SCA10), which is caused by an ATTCT pentanucleotide expansion within intron 9 (Matsuura et al., 2000). The consequences of the pentanucleotide expansion on Atxn10 function or non-expansion coding mutations on the function of ATXN10 remain unclear. Investigation of the pentanucleotide expansion mutation indicates that the allele is transcribed at normal levels and is spliced normally (Wakamiya et al., 2006). To date, the only reported human incidence of ATXN10 mutation (IVS8-3T>G) was observed in three Turkish siblings from a consanguineous family. This mutation resulted in Nephronophthisis-like kidney defects that ultimately led to death as infants (Sang et al., 2011). This same study further identified ATXN10 as a Nephronophthisis (NPHP) and Joubert Syndrome (JBTS) associated gene that indirectly interacts with the ciliary transition zone protein, NPHP5, near the base of the cilium. NPHP is a form of medullary cystic kidney disease with associated nephron loss (Luo and Tao, 2018), while JBTS is autosomal recessive or X-linked cerebellar ataxia associated with cerebellar hypoplasia (Romani et al., 2013). Both NPHP and JBTS fall into the class of disorders collectively termed ciliopathies. Ciliopathies result from improper structure or function of the primary cilium. These small microtubule based appendages are present on the surface of nearly every mammalian cell type and are crucial for mediating many cell signaling events (Sharma et al., 2008).

Knockdown of *Atxn10* in rat primary cortical and especially cerebellar neurons is cytotoxic (Marz et al., 2004). Interestingly, overexpression of *Atxn10* alone is sufficient to induce neuritogenesis in neuronal precursor cells where it interacts with the G-protein β2 subunit to drive activation of the RAS-MAPK-ELK-1 signaling cascade (Waragai et al., 2006). Furthermore, Aurora B phosphorylation of ATXN10 promotes its interaction with polokinase 1 (Plk1) (Tian et al., 2015). This interaction between ATXN10 and Plk1 is necessary for cytokinesis *in vitro* (Li et al., 2011). The function of ATXN10 *in vivo* remains largely unresolved.

To initiate studies into the *in vivo* functions of ATXN10, we established congenital (*Atxn10*^*KO*^) and conditional (*Atxn10*^*flox*^) mutant mice and assessed the consequence of ATXN10 loss during both embryogenesis and in adult tissues. Congenital loss of ATXN10 results in severe cardiac development abnormalities and gestational lethality. Tissue specific ablation of ATXN10 in the developing endothelium and myocardium similarly result in embryonic lethality. Induction of ATXN10 loss in adult mice causes lethality due to moderate to severe pancreatic, renal, and gastrointestinal abnormalities, and severe defects in glucose homeostasis. Further analysis of renal phenotypes revealed an epithelial to mesenchymal transition (EMT) of the kidney tubule epithelial cells. Similarly, in the pancreas, acinar cells appear to undergo a transdifferentiation process resulting in more progenitor-like phenotypes.

Previous work indicates that ATXN10 is predominantly a cytoplasmic protein with cell cycle dependent localization of the phosphorylated protein (on Serine 12), to the Golgi during interphase, the centrioles during prophase, and the midbody during telophase. Our studies similarly indicate that localization of ATXN10 is predominantly cytoplasmic; however, it is enriched near the centrioles and at the base of the primary cilium. While a ciliary role for ATXN10 cannot be excluded, we show that loss of ATXN10 does not affect ciliogenesis in fibroblast or epithelial cells, although acini in *Atxn10* postnatal-induced mutants do exhibit ectopic cilia possibly associated with changes in cell type.

## Results

### Loss of ATXN10 does not affect cilia formation or maintenance

To determine the localization of ATXN10, we generated an EGFP tagged ATXN10 (ATXN10::EGFP) for expression in cultured cells. Overexpression of ATXN10::EGFP in Inner Medullary Collecting Duct (IMCD) cells supports a predominantly cytoplasmic expression pattern; however, enrichment of ATXN10::EGFP near Fgfr1op (FOP) positive centrioles or basal bodies is seen in 79.3% of transfected cells, regardless of whether they had a primary cilium. Of the transfected cells that have cilia, we detected an enrichment of ATXN10 around the base of all cilia (**Figure 1A).** This information led to the investigation of whether ATXN10 is necessary for ciliogenesis. In Mouse Embryonic Fibroblasts (MEFs) generated from *Atxn10*^*KO*^ embryos, there was a trend toward fewer cilia, but these differences were not statistically significant between *Atxn10*^*KO*^ (33.5%) and control (59.8%) cells (p=0.07) (**Figure 1B**). Similar to published reports, MEFs generated from *Atxn10*^*KO*^ embryos display cell cycle abnormalities creating difficulties in maintaining the cells for longer than two or three passages (see below). As the formation of the primary cilium is tied to the cell cycle (Malicki and Johnson, 2017), we wanted to determine whether the loss of ATXN10 affected ciliary maintenance following cilia formation. To address this question, we generated conditional MEFs using *Atxn10*^*Cagg*^ embryos, induced ATXN10 loss prior to or after confluency and then serum starved to induce cilia formation. If induction occurred prior to confluency, *Atxn10*^*Cagg*^ MEFs (35.1%) exhibited a non-significant trend towards fewer cilia compared to controls (42.7%) (p=0.42) (**Figure 1C**). Induction of *Atxn10*^*Cagg*^ MEFs post-confluency resulted in similar percentages of ciliated cells between control (76.3%) and *Atxn10*^*Cagg*^ MEFs (72.0%) (p=0.51) (**Figure 1D**).

**Figure 1:**
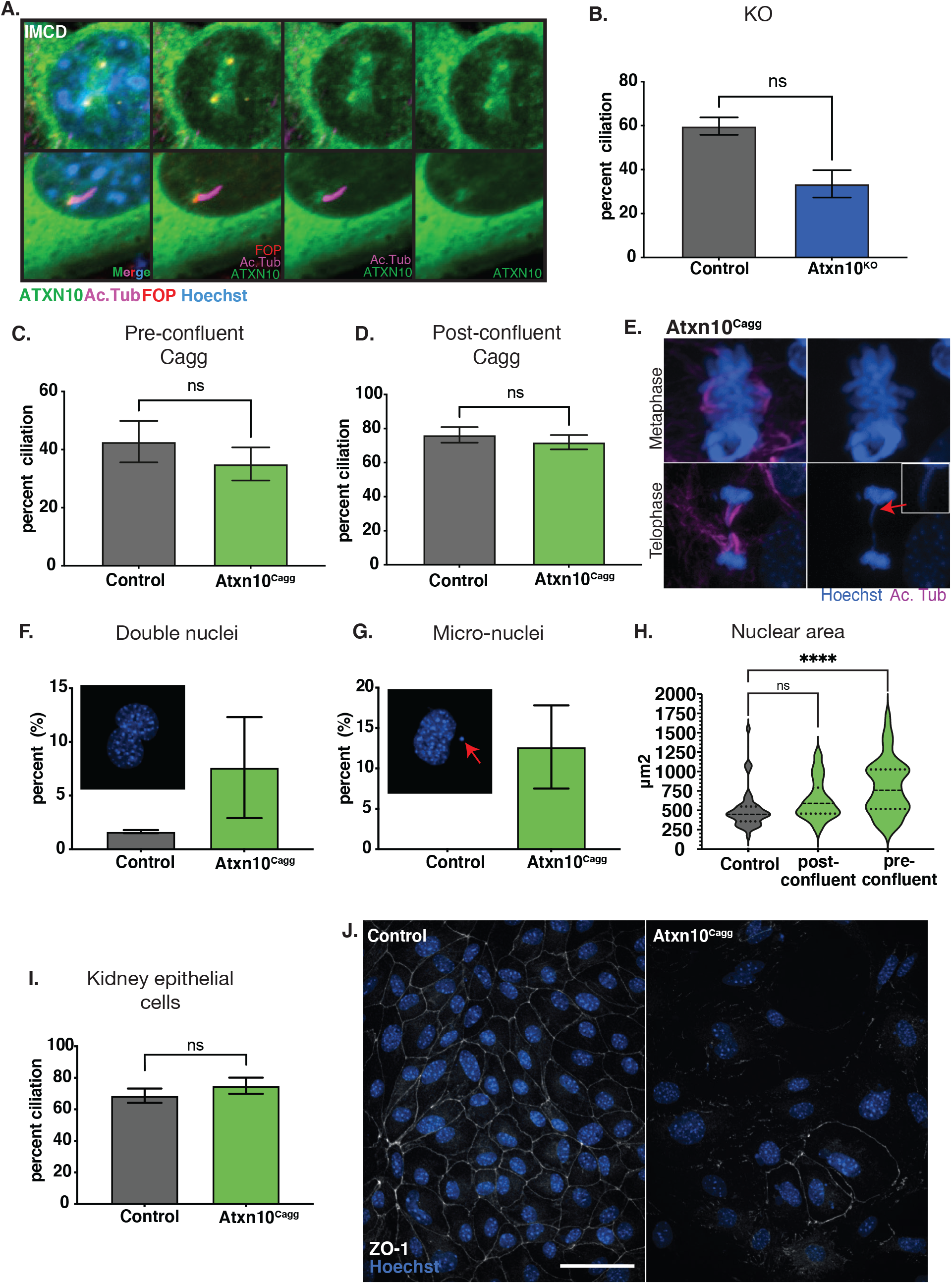
*In vitro* analysis. A) ATXN10::EGFP localized to the cilia basal body and centrioles in IMCD cells stained for cilia, Acetylated α-tubulin (Ac. Tub, purple) and basal bodies, FGFR1 oncogene partner (FOP, red), and hoechst (blue) for nuclei. B) Percent ciliation in control and *Atxn10*^*KO*^ MEFs. C) Percent cilia in control and *Atxn10*^*Cagg*^ MEFs when Cre is induced prior to confluency. D) Percent cilia in control and *Atxn10*^*Cagg*^ MEFs when Cre is induced after confluency. E) Nuclear abnormalities observed in metaphase and telophase of the cell cycle in *Atxn10*^*Cagg*^ MEFs. F and G) Percent of primary kidney epithelial cells containing two nuclei and exhibiting nuclear blebbing and micro-nuclei formation. H) Nuclear area (μm^2^) in primary kidney epithelial cells in control, and induced post and pre-confluent cells. I) Percent cilia in primary kidney epithelial cells from control and *Atxn10*^*Cagg*^ animals that were post-confluency. J) Immunofluorescence staining for tight junction protein, zonula occludens-1 (ZO-1, white) in non-induced (control) and induced mutant (*Atxn10*^*Cagg*^) primary kidney epithelial cells (scale bar= 50μm). Statistical significance of nuclear area was determined using a one-way ANOVA with multiple comparisons. All other statistical significance was determined using unpaired T-test.

Although cilia formation in MEFs is not significantly affected by loss of ATXN10 in cultured cells, we did observe abnormalities in cell division based on the frequency of irregular spindle formations observed along with chromosomal bridges (**Figure 1E, red arrow**). To observe the effect of ATXN10 loss in an epithelial cell line, primary kidney epithelial cells were isolated from *Atxn10*^*Cagg*^ mice. Similar to what was observed in MEFs, loss of ATXN10 prior to confluency resulted in cells that failed to grow to confluency and could not be maintained (data not shown). *Atxn10*^*Cagg*^ primary renal epithelial cells displayed an increased prevalence of cells with two nuclei (7.6% in *Atxn10*^*Cagg*^ compared to 1.7% in controls, (p=0.33)) (**Figure 1F**). They also exhibited a large increase in nuclear blebbing and micro nuclei formation (12.7% in *Atxn10*^*Cagg*^ compared to 0% in controls) (**Figure 1G**) and nuclear size was increased in cells induced prior to confluency (p<0.0001; with nuclear area in control cells averaging 504μm^2^, pre-confluent induced cells averaging 632μm^2^, and post-confluent induced averaging 799 μm^2^) (**Figure 1H**). The frequency with which post-confluent induced primary kidney epithelial cells presented a cilium was not different between control and *Atxn10*^*Cagg*^ cells (68.6% in control cells versus 74.9% in *Atxn10*^*Cagg*^ cells) (**Figure 1I**). Another consistent observation regarding the primary kidney epithelial cells was an increase in cell spreading and more fibroblast-like cell morphology following Cre induction (**Supplemental figure 1F, Figure 1J**). Staining for the epithelial tight junction marker ZO-1 shows a distinct loss of localization in *Atxn10*^*Cagg*^ primary kidney epithelial cells compared to non-induced controls (**Figure 1J**). Collectively, this supports ATXN10’s role in cell division and that it transiently accumulates around the ciliary basal bodies; however, it is dispensable for ciliogenesis. These data also indicate a potential role for ATXN10 in maintaining epithelial cell phenotypes like tight junctions.

### Atxn10 In Vivo expression analysis

The promoter driven expression of *LacZ* in the Tm1a knockout first (KO) allele (**Figure 2A)** allowed gene expression to be examined via β-galactosidase staining. Staining performed on heterozygous *Atxn10*^*KO/+*^ and wildtype control embryos indicate that *Atxn10* expression begins in the developing heart tube at embryonic day 8.5 (E8.5) and is predominantly confined to the developing heart with expression beginning to expand to the surrounding mesoderm and parts of the neural tube at E10.5 (**Figure 2B and C**). By E11.5 reporter expression has expanded to the entire embryo. β-galactosidase staining performed on sections of E10.5 *Atxn10*^*KO/+*^ embryos shows expression of *Atxn10* in both the myocardium and endocardium (**Figure 2C**).

**Figure 2:**
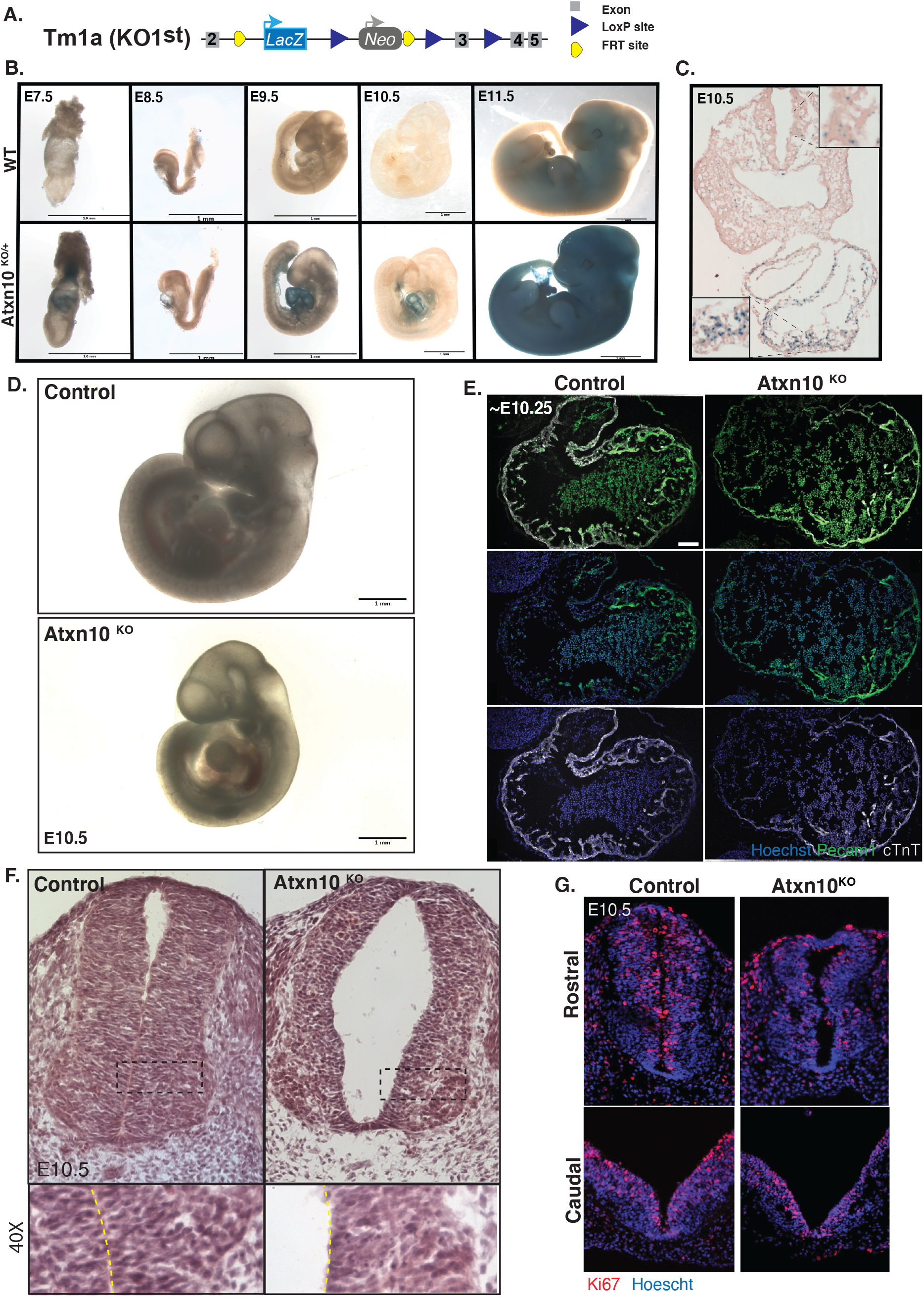
Phenotypic analysis of congenital loss of ATXN10. A) Schematic depicting the Atxn10 KO1st allele. B) β-Galactosidase staining in wild-type (top) and heterozygous (bottom) animals at E7.5, E8.5, E9.5, E10.5, and E11.5. C) β-Galactosidase staining of a crossection of an E10.5 heterozygous embryo. D) Images of wild-type (top) and *Atxn10*^*KO*^ (bottom) embryos at E10.5 (scale bar = 1mm) E) Immunofluorescence staining for PECAM1 (green), cTnT (white), and hoechst (blue) in the heart of wild-type and *Atxn10*^*KO*^ embryos at E10.5 (scale bar= 100 μm). F) H&E staining of control and *Atxn10*^*KO*^ rostral neural tubes at E10.5. G) Ki67 and nuclei staining of Caudal (top) and rostral (bottom) neural tubes in WT and Atxn10^KO^mutant animals.

### Congenital loss of ATXN10 results in pericardial effusion and embryonic lethality

In agreement with the embryonic lethality reported by the International Mouse Phenotyping Consortium (IMPC), we found that Atxn10 mutant embryos die shortly after E10.5 (Wakamiya et al., 2006;Dickinson et al., 2016). Embryos at E10.5 are generally smaller than controls and begin to show pericardial edema (**Figure 2D**). By E10.5 severe pericardial effusion is observed with lethality occurring shortly after (**Figure 2D**). Immunofluorescence staining of sections through the heart region using markers for the endothelium (PECAM1) and the myocardium (cTnT) show that while both layers are present in *Atxn10*^*KO*^ embryos, the walls of the developing heart are thinned and the heart is larger in volume (**Figure 2E**).

### Congenital loss of ATXN10 results in neural tube defects

In addition to the cardiac defects mentioned above, structural abnormalities were also observed in the neural tubes of mutant embryos (**Figure 2F and G**). H&E staining of E10.5 embryos highlight the thin, disorganized structure of the neural epithelium (**Figure 2F**) and although the rostral neural tubes have closed, they exhibit an increase in the luminal space. While the caudal neural tube has not fully closed in the control nor mutant embryos, Ki67 staining in both rostral and caudal neural tubes indicate that the thinned appearance of the neural tube is likely not due to a lack of proliferation (**Figure 2G**).

### Tissue specific ablation of ATXN10 in the myocardium and endothelium results in embryonic lethality

At the stage that pericardial effusion is observed in *Atxn10*^*KO*^ embryos, interactions between the developing endocardium and myocardium are crucial for proper development (Samsa et al., 2013). To determine if the cardiac phenotype seen in congenital knockout mice is specifically due to loss of ATXN10 in the developing myocardium or endocardium, the conditional *Atxn10* allele was used (**Figure 3A**). By inducing loss of ATXN10 in the developing endothelium using the Tek2-Cre (Tie2) (Koni et al., 2001), embryos exhibit lethality between E11.5 - E13.5. *Atxn10*^*Tie2*^ embryos collected at E10.5 are grossly indistinguishable from control littermates (**Supplemental figure 2A**). Embryos isolated at E12.5 can be distinguished from littermates due their pale coloration and pooling of blood around the heart (**Figure 3B**). Embryos that can still be recovered at E13.5 show severe vascular abnormalities (**Supplemental figure 2B**). Staining of *Atxn10*^*Tie2*^ mutants prior to death indicate that the endothelium is still present in both the developing heart (**Figure 3C and 3D**) and in the embryonic mesoderm (**Figure 3E**).

**Figure 3:**
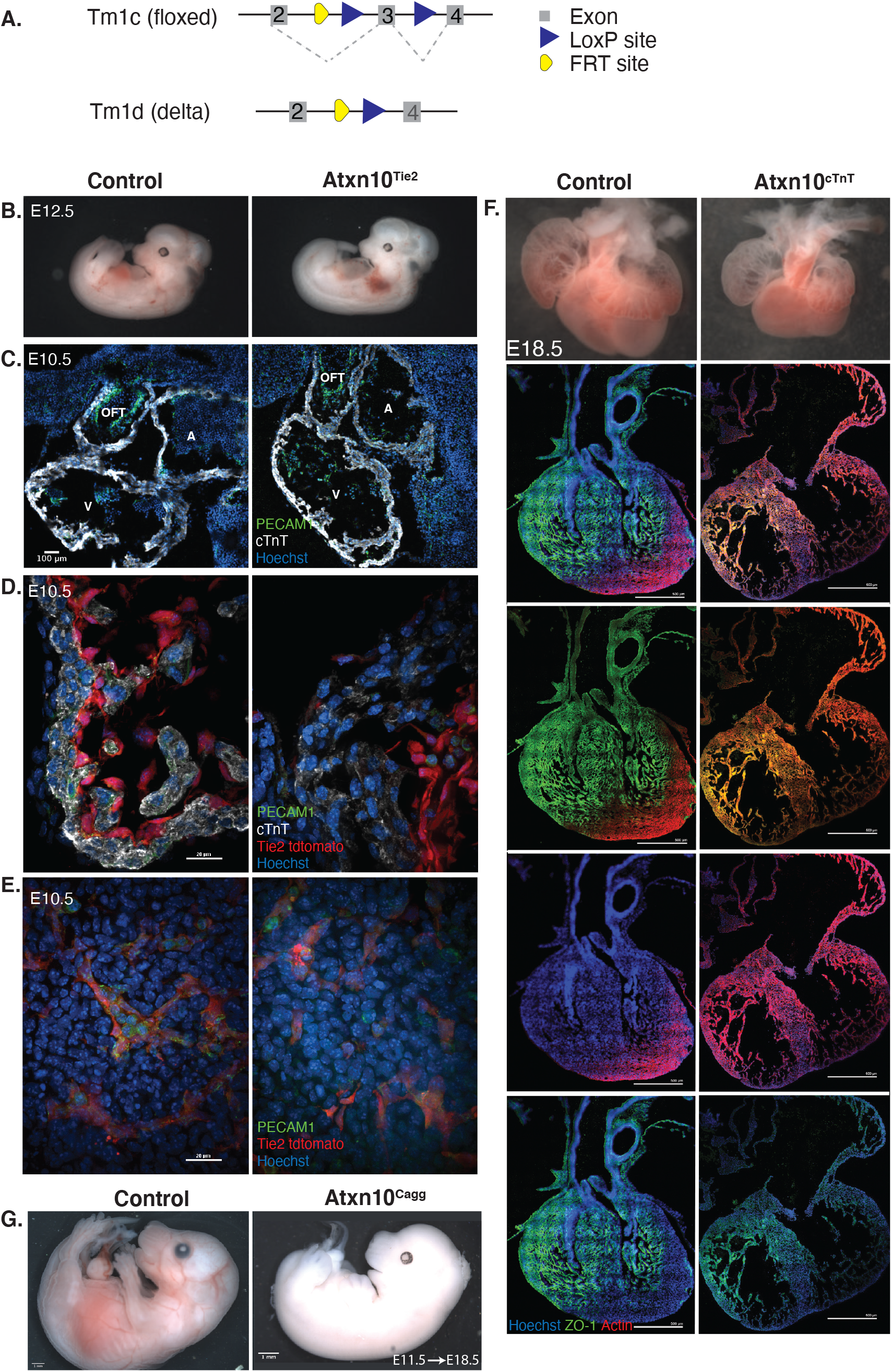
Tissue specific ablation of ATXN10. A) Schematic depicting the *Atxn10* tm1c (floxed) allele and schematic depicting the *Atxn10* tm1d (delta) allele. B) Control and *Atxn10*^*Tie2*^ mutant embryos at E12.5. C) Immunofluorescence staining of cardiac crossections taken from E10.5 control and *Atxn10*^*Tie2*^ embryos with markers for PECAM1 (green), cTnT (white), and hoechst (blue) (scale bar= 100 μm.) D) Immunofluorescence staining of cardiac trabeculae in E10.5 control and *Atxn10*^*Tie2*^ embryos with green representing PECAM1, white representing cTnT, and nuclei shown in blue. Activity of Tie2-Cre is marked in red by tdtomato reporter (scale bar= 100 μm). E) Immunofluorescence staining of vasculature in E10.5 mesoderm in control and *Atxn10*^*Tie2*^ embryos with PECAM1 (green), cTnT (white), hoechst (blue), and Tie2-Cre activity is marked in red by tdtomato reporter (scale bar= 100 μm). F) Hearts isolated from control and *Atxn10*^*cTnT*^ embryos at E18.5. Immunofluorescence staining indicates ZO-1 shown in green, actin shown in red, and nuclei (blue) (scale bar= 500 μm). G) images of Control and *Atxn*10^Cagg^ embryos induced in utero at E11.5 and isolated at E18.5 (scale= 1mm).

Using a cTnT-Cre we also assessed the effect of ATXN10 loss in the myocardium (Jiao et al., 2003). Ultimately these mutants are still embryonic lethal during late gestation; however, at E18.5 *Atxn10*^*cTnT*^ embryos appear relatively normal except for mild edema around the neck region of the embryo (data not shown). Closer observation of the heart indicates reduced trabeculation and ventricular non-compaction compared to controls (**Figure 3F**).

The spatiotemporal specificity of gene expression and the failure of the two tissue-specific mutants to phenocopy the congenital heart phenotype while still resulting in lethality led to the question of whether embryonic lethality is specific to cardiac abnormalities. To test this, timed matings were established between *Atxn10*^*flox/flox*^ females and *Atxn10*^*flox/flox*^; *Cagg-Cre* positive males. Pregnant females were then induced 11.5 days into pregnancy. Seven (7) days following induction (E18.5), the resulting Cre positive embryos exhibited a reduction in the presence of blood throughout the embryo and their hearts were no longer beating indicating they were nonviable (**Figure 3G**). Although these embryos had been induced following the initial cardiac morphological events that were impaired in *Atxn10*^*KO*^ embryos, lethality was still ultimately the outcome supporting the conclusion that lethality is due to additional defects outside the heart and blood vessels.

### Loss of ATXN10 in adult mice results in pancreatic, renal, and gastrointestinal abnormalities followed by abrupt lethality

To determine whether ATXN10 is necessary postnatally, loss of ATXN10 was induced in *Atxn10*^*Cagg*^ adult mice at 4 weeks and 8 weeks of age. Surprisingly, this had dramatic effects with mice induced at 4 weeks of age failing to gain weight (**Supplemental figure 3A**). When animals were induced at 8 weeks of age only male mice exhibited a significant weight difference at 17 days post-induction while female mice seemed to maintain a weight comparable to controls (**Figure 4A**). Strikingly, *Atxn10*^*Cagg*^ (Cre positive) mice induced at 4 weeks (data not shown) and 8 weeks of age resulted in abrupt lethality between 16-26 days post-induction (**Figure 4B**). Pathological analysis of tissues isolated from control and *Atxn10*^*Cagg*^ animals prior to death indicates that a major contributor to their death is most likely a result of pancreatic acinar injury. Histology of *Atxn10*^*Cagg*^ pancreata show cytoplasmic basophilia and acinar cell necrosis with significant lymphocytic infiltrates within and around affected acini indicating a recent injury to the pancreas (**Figure 4C**). Analysis of serum glucose levels in non-fasted animals show a significant decrease in blood glucose levels in *Atxn10*^*Cagg*^ animals compared to control (**Figure 4D**). Compared to control, the pancreata in *Atxn10*^*Cagg*^ animals are significantly smaller in size (**Figure 4E**). Lethality is likely due to the combined effect of pancreatic damage and reduced food intake indicated by hyperkeratosis of the non-glandular region of the gastrointestinal tract and hepatocyte atrophy (Supplemental figure 3C). Further analysis of the stomach in *Atxn10*^*Cagg*^ animals revealed moderate lymphocytic and neutrophilic infiltrate of the submucosal and mucosal regions. Additionally, mild crypt dilation, epithelial necrosis, and diffuse, mild epithelial hyperplasia, dysplasia, and mild mineralization indicated chronic, active gastritis and secondary epithelial changes (data not shown). Interestingly, these indicators of an ongoing insult to the glandular epithelium were not observed in the non-glandular regions of the gastrointestinal tract (duodenum/jejunum, cecum/colon, omentum/mesentery).

**Figure 4.**
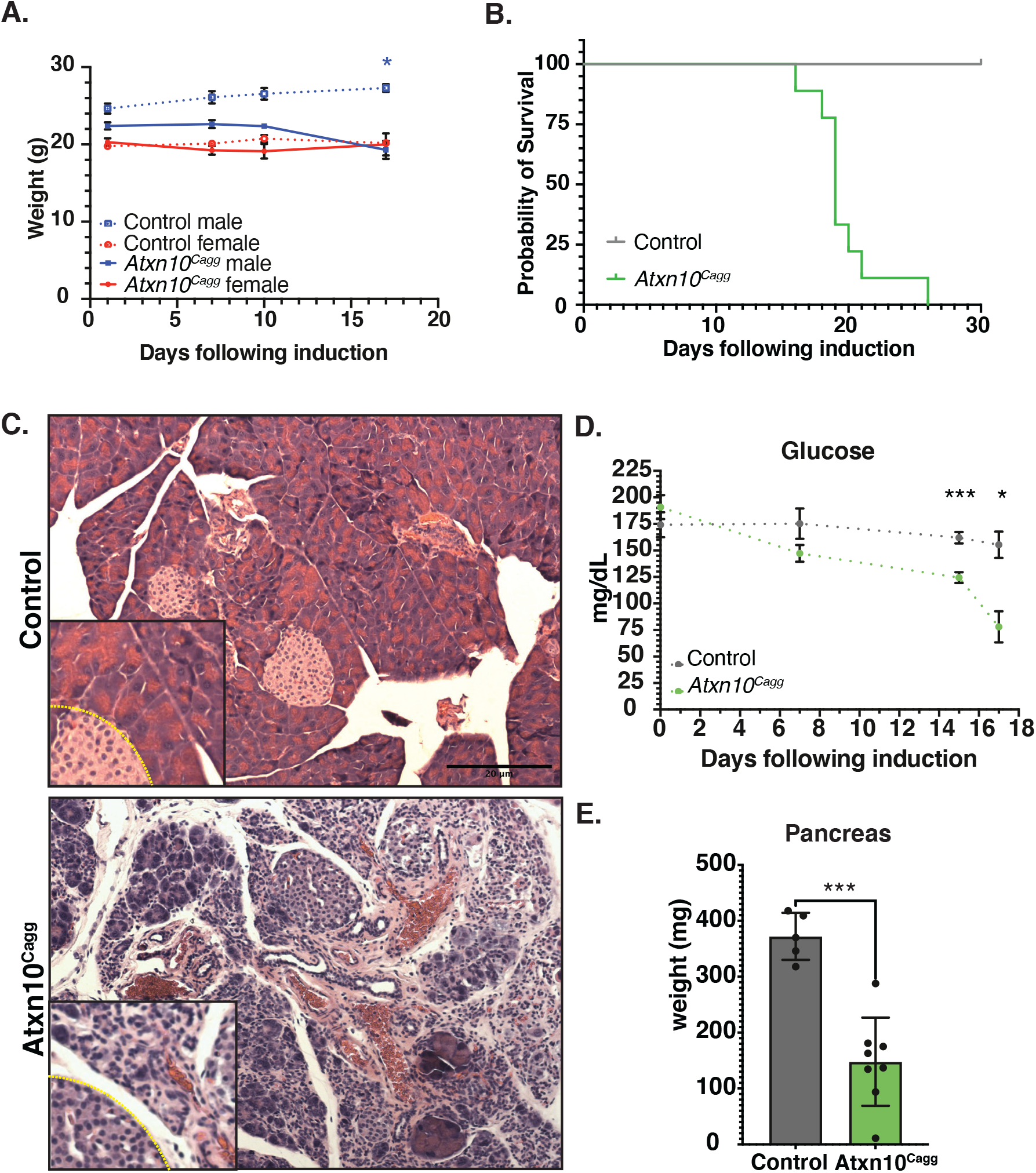
Pancreas defects. A) Weights following induction of Control and *Atxn*10^Cagg^ animals at 8 weeks old (P= 0.468 in male mice at 17 days post induction; N= 7 Cre-females, 2 Cre-males, 6 Cre+ females, and 4 Cre+ males). B) Kaplan-meier survival curve of Control (gray) and *Atxn*10^Cagg^ (green) animals induced at 6 weeks old (P<0.0001; N=9 Cre- and 9 Cre+ animals). C) H&E staining of *Atxn10*^*flox/flox*^ (left) and *Atxn*10^Cagg^ (right) pancreas. D) Levels of blood glucose in nonfasted mice (P=0.002 at 15 days post induction and P=0.0399 at 17 days post induction; N=5 Cre- and 8 Cre+ up to 15 days post induction and N= 3 Cre- and 4 Cre+ mice at day 17). E) Pancreatic weight at 15–17 days post induction (P=0.0001, N=5 Cre- and 8 Cre+ animals). Statistical significance was determined using mixed effects analysis with multiple comparisons for change in weight and change in glucose levels over time. Statistical significance of survival was determined via log-rank test. Statistical significance of pancreas weight was determined using unpaired T-test.

While pathological analyses of the retina, liver, lung, and spleen (**Supplemental figure 3B-3E**) were unremarkable, the kidney presented with histological markers suggestive of a regenerative response to acute tubular injury marked by cytoplasmic basophilia, an increased nuclear to cytoplasmic ratio, open chromatin, distinct nucleoli and occasional mitotic figures in the proximal tubules (**Figure 5A**). Despite these histological findings, blood serum analysis revealed that while alkaline phosphatase (**Figure 5B**) and blood urea nitrogen (BUN) (**Figure 5C**) levels are increased in *Atxn10*^*Cagg*^ animals, serum albumin (**Figure 5D**), sodium (**Figure 5E**), calcium (**Figure 5F**), phosphorus (**Figure 5G**), creatinine (**Figure 5H**), and total protein (**Figure 5I**) are not significantly different between control and *Atxn10*^*Cagg*^ animals.

**Figure 5.**
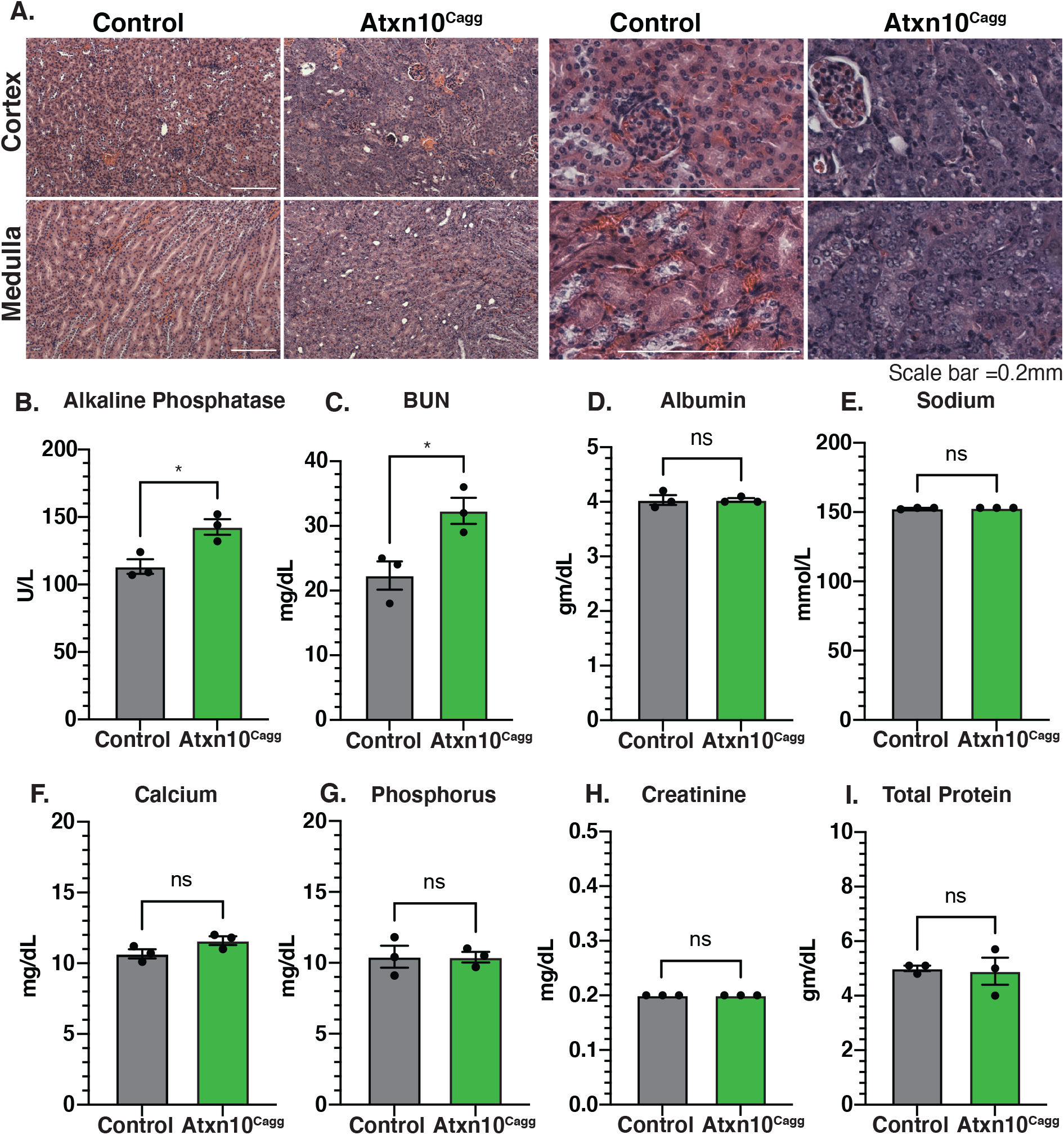
Renal defects. A) H&E staining of Control (left) and *Atxn*10^Cagg^ (right) kidneys 17 days post induction. Blood serum levels of B) Alkaline Phosphatase (P= 0.02), C) BUN (P=0.03), D) Albumin, E) Sodium, F) Calcium, G) Phosphorous, H) Creatinine, and I) total protein in Control and *Atxn*10^Cagg^ animals 17 days post induction. Scale bars=50μm. Statistical analysis was performed using unpaired T-test.

### Loss of ATXN10 causes the pancreatic epithelium to become more progenitor-like and results in ectopic primary cilia growth

Within the pancreas, acinar cells can be identified by the presence of amylase while ductal cells express SOX9 (Kopp et al., 2012) In control pancreata, ductal cells can be identified by SOX9 positive nuclei that are frequently also positive for the proliferation marker Ki67. In *Atxn10*^*Cagg*^ pancreata there are a large number of cells that are positive for SOX9 and Ki67 but lack amylase, suggesting an increase in the number of ductal cells. Intriguingly, there are also cells that are positive for SOX9 and Ki67 that also express amylase. In the most severely affected pancreata, a population of ciliated cells that lack SOX9, amylase, and Ki67 are present (**Figure 6A**). Closer evaluation of *Atxn10*^*Cagg*^ pancreata showed a substantial increase in Vimentin positive cells accompanied by a decrease in E-Cadherin expression. Cells that did maintain E-Cadherin expression exhibited highly disorganized E-Cadherin cellular localization (**Figure 6B**). Compared to control pancreata, *Atxn10*^*Cagg*^ pancreata exhibit a significantly higher number of Ki67 positive cells (P=0.0004; N= 3 Cre- and 4 Cre+ animals) (**Figure 6C and D**). Furthermore, *Atxn10*^*Cagg*^ pancreata appear to exhibit an increase in the density of primary cilia in the exocrine region of the pancreas (**Figure 6E**). Arl13b positive cilia are normally present throughout the islet of the control pancreas, but are infrequent in the exocrine region (**Supplemental figure 5, left**). In *Atxn10*^*Cagg*^ pancreata the islets exhibit longer cilia, and the exocrine regions of the pancreas, which are typically nonciliated, have a high density of ciliated cells (**Supplemental figure 5, right**). Collectively this indicates that pancreatic cells are becoming more progenitor or mesenchymal-like.

**Figure 6.**
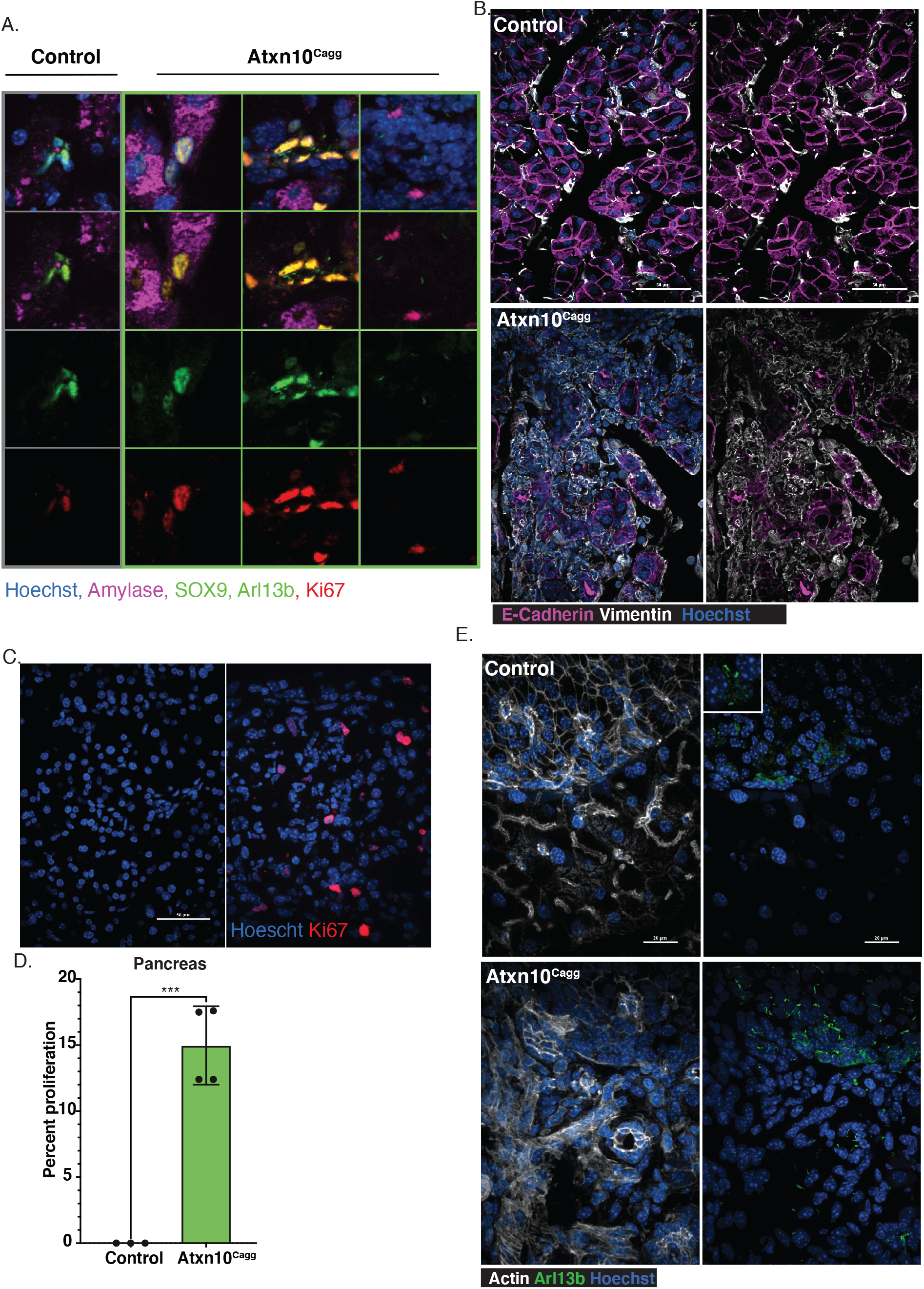
Loss of epithelial characteristics in adult *Atxn10*^*Cagg*^ pancreata. A) Immunofluorescence staining for amylase (purple), SOX9 (green), Arl13b (green), Ki67 (red), and Hoechst (blue) in control and *Atxn*10^Cagg^ pancreata. B) Immunofluorescence staining for E-cadherin (purple), Vimentin (white), and Hoechst (blue) in Control (top) and *Atxn10*^*Cagg*^ (bottom) pancreas. C) Images and D) quantification of proliferation in the pancreas (P=0.003; N=3 Cre- and 4 Cre+ animals) as shown by Ki67 staining (red) (scale bar= 50 μm). E) Immunofluorescence staining for Actin (white), cilia shown by Arl13b staining (green) and Hoechst (blue) in Control (left) and *Atxn*10^Cagg^ (right) pancreata (scale bar= 20 μm).

### Loss of ATXN10 induces proliferation and structural abnormalities in the kidney

Analysis of the kidney by immunofluorescence staining for LTA (proximal tubules) and DBA (collecting tubules/ducts) identified tubule segments in *Atxn10*^*Cagg*^ animals in which LTA and DBA colocalize, and in many tubules LTA is no longer restricted to the apical surface of the cells (**Figure 7A**). Actin staining in *Atxn10*^*Cagg*^ kidneys further highlights structural disorganization. Actin organization in control kidneys shows the normal dense actin staining on the apical side of the renal epithelium compared to basolateral edges of the cell. In *Atxn10*^*Cagg*^ kidneys this staining is no longer enriched at the apical surface but rather, there is actin accumulation around the entire cellular membrane in many tubules (**Figure 7B, red**). Furthermore, staining for the presence of cilia with acetylated α-tubulin shows disorganized punctate cilia (**Figure 7B, green**). To further support the loss of tubular structure in *Atxn10*^*Cagg*^ kidneys, we examined localization of the tight junction marker ZO-1. *Atxn10*^*Cagg*^ samples show a disorganized appearance of ZO-1 staining in the cytoplasm compared to the expected membrane associated staining indicative of mature tight junctions (**Figure 7B, white**). In summary, the kidneys exhibit characteristics of disrupted epithelial polarity such as loss of apical localization of LTA, F-actin remodeling, small punctate cilia, and loss of ZO-1 localization at the cell-to-cell contacts. Furthermore, compared to control kidneys, *Atxn10*^*Cagg*^ kidneys exhibit a significantly higher number of Ki67 positive cells (P=0.003; N= 3 Cre- and 4 Cre+ animals) (**Figure 7C and D**).

**Figure 7.**
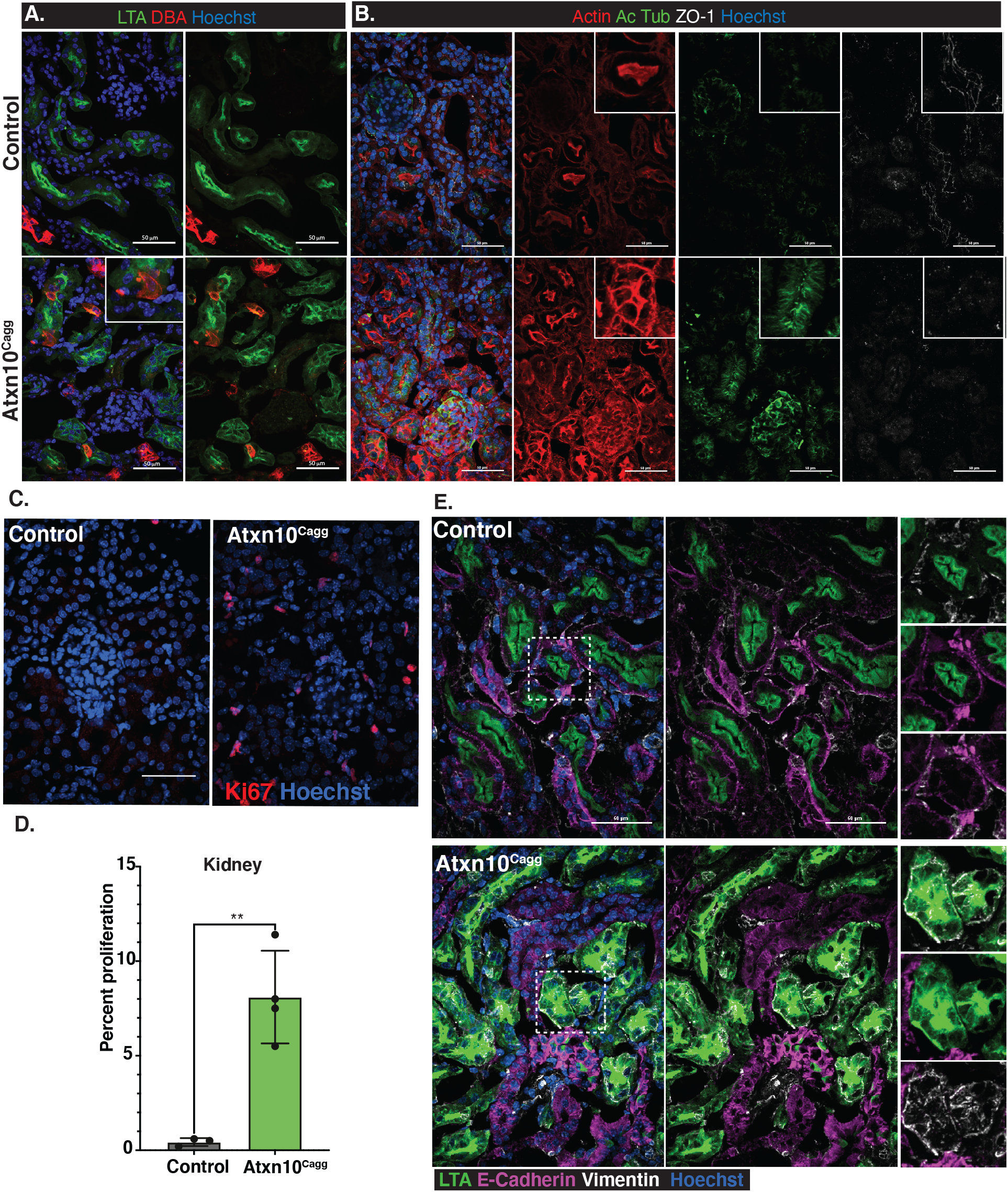
Loss of epithelial characteristics in adult induced *Atxn10*^*Cagg*^ kidney. A) Staining for proximal tubule (LTA, green) and collecting duct (DBA, red) in Control (top) and *Atxn*10^Cagg^ (bottom). B) Staining for Actin (Phalloidin, red), cilia (Acetylated α-tubulin, green), and ZO-1 (white) in Control (top) and *Atxn*10^Cagg^ (bottom) kidneys. C) Images and D) quantification of proliferation in the kidney (P=0.0004; N=3 Cre- and 4 Cre+ animals) as shown by Ki67 (red) staining. E) Immunofluorescence staining for LTA (green), E-cadherin (purple), Vimentin (white), and Hoechst (blue) in Control (top) and *Atxn10*^*Tie2*^ (bottom) kidneys. (scale bar=50 μm)

### Loss of ATXN10 results in more mesenchymal-like cells in renal tubules

Collectively, the mixed tubule identity paired with loss of apical restriction of LTA, actin remodeling, and loss of ZO-1 at the cell membrane in the renal proximal tubules led to the question of whether epithelial cells were becoming more mesenchymal-like. To test this, Cre negative induced (control) and Cre positive (*Atxn10*^*Cagg*^) induced kidneys were stained for E-Cadherin and Vimentin to identify epithelial and mesenchymal cells, respectively. In control kidneys, E-Cadherin staining is observed at the basolateral membrane and at areas of cell-to-cell contact throughout the tubules. Vimentin staining is adjacent to the E-Cadherin positive basolateral membrane of the renal tubules. In contrast, cells in *Atxn10*^*Cagg*^ tubules show LTA positive staining that is spread diffusely throughout the cell. Furthermore, these LTA positive cells do not express E-Cadherin, but rather become positive for the mesenchymal marker, Vimentin (**Figure 7E**).

## Discussion

Previous studies of *Atxn10* have focused on its role in SCA10, a pentanucleotide expansion disorder (Matsuura et al., 2000), but there is limited analysis of the direct consequence of mutations affecting coding regions. Studies highlighting the role of ATXN10 in cell biology have been performed *in vitro* and centered on its role in cell division. Additionally, studies using the G-LAP-Flp purification strategy in Intermedullary Collecting Duct Cells (IMCDs) identify ATXN10 as having indirect interactions with the ciliary transition zone and ciliopathy protein NPHP5 (Torres et al., 2009;Sang et al., 2011), suggesting that ATXN10 may play a role in cilia function or formation. Our findings show that ciliogenesis is not affected upon the loss of ATXN10 *in vitro* or *in vivo*. Consistent with previous reports, ectopic expression of ATXN10 in cultured cells shows diffuse localization within the cytoplasm with enrichment at the centrioles and base of the cilium. Previously reported interactions between ATXN10 and NPHP5 further support that ATXN10 is localized at the base of the cilium. Previous work has shown cell cycle specific localization of phosphorylated ATXN10. In cultured *Atxn10*^*Cagg*^ renal epithelial cells grown to confluency and then induced for Atxn10 loss, cells adopt a more fibroblast-like appearance, and the localization of ZO-1 to the sites of cell-to-cell contact is disrupted. In addition, we observe defects in chromosomal segregation with a high frequency of micronuclei along with chromosomal bridges. As a consequence, it is difficult to maintain cells in culture once Atxn10 loss is induced.

*In vivo,* we see that Atxn10 function is essential for viability. Expression of *Atxn10* is restricted to the developing heart until after E10.5 when expression expands rapidly to cells throughout the embryo. Not surprisingly, based on the highly localized expression pattern in the early embryo, loss of Atxn10 resulted in severe pericardial effusion and ultimately cardiac failure. While *Atxn10* expression is concentrated in the developing heart through E10.5, defects are observed in the epithelial cells of the neural tube in *Atxn10*^*KO*^ embryos. This is the earliest indication that ATXN10 plays a role in epithelial and endothelial cell maintenance.

To determine if ATXN10 is required in a tissue/cell type specific manner in the cardiovascular system during embryogenesis, *Atxn10* was deleted in the developing myocardium and endothelium using cTnT-Cre and Tie2-Cre, respectively. Both of these tissue specific conditional *Atxn10* knockouts result in embryonic lethality. In the case of *Atxn10*^*Tie2*^ embryos, lethality typically occurs between E11.5-E13.5. In *Atxn10*^*Tie2*^ embryos obtained at E10.5, the developing vasculature is present indicating that lethality is not a result of a failure in vasculogenesis; however, Tie-2 transgene activity is also present in hematopoetic progenitor cells, and this may also contribute to lethality in these animals (Tang et al., 2010)

Comparatively, *Atxn10*^*cTnT*^ embryos exhibit cardiac abnormalities similar to those seen in null embryos but survive longer than *Atxn10*^*KO*^ or *Atxn10*^*Tie2*^ embryos. This could be due in part to delayed activation of Cre in this line or to mosaicism in expression of the Cre. Regardless, at E18.5 the hearts of *Atxn10*^*cTnT*^ embryos display reduced ventricular wall thickness and disorganization of ZO-1 and actin reminiscent of the defects observed in *Atxn10*^*KO*^ embryos at earlier time points.

Interestingly, the importance of ATXN10 is not limited to embryonic development. Using the inducible Cagg-Cre model, loss of ATXN10 in adult animals causes a rapid decline in health and results in lethality approximately 3 weeks post-induction. Necropsy and histological analysis point to moderate to severe pancreatic abnormalities and gastritis paired with reduced food intake as the leading cause of lethality. Additional renal abnormalities likely also contribute to their progressively worsening condition. Furthermore, an increase in purple staining, or cytoplasmic basophilia, was observed in *Atxn10*^*Cagg*^ pancreata and kidneys. This phenomena is a result of RNA in the cytoplasm and is indicative of a regenerative epithelial cell or a precursor state in which a cell has recently stopped dividing (Chan, 2014).

EMT is a vital process during development. Post-developmentally it is associated with cancer metastasis, migration, and invasion (Yamashita et al., 2018). The cellular events included in this process are disruption to cell polarity, loss of epithelial cell-to-cell junctions, downregulation of epithelial markers such as E-cadherin and Zonula occludens (ZO-1) paired with an up-regulation of mesenchymal markers such as Vimentin, alterations to the cytoskeletal architecture, and an increase in secretory abilities. *Atxn10*^*Cagg*^ kidneys exhibit four of these characteristics: loss of apical restriction of LTA, dissolution of ZO-1 localization at the membrane, downregulation of E-cadherin paired with an up-regulation of Vimentin, and alterations in the actin cytoskeletal network. Collectively, these point to EMT in *Atxn10*^*Cagg*^ kidneys. EMT of renal tubule epithelial cells is associated with the kidney’s injury and repair process (Humphreys et al., 2008). Further indication that the kidney is attempting to repair following an injury is an increase in Ki67 positive cells. In the kidney, baseline proliferation is normally very low. While Ki67 is commonly used as a proliferative marker, cells begin to acquire Ki67 in the nucleus during S phase of the cell cycle, and its presence persists throughout the G2 and M phases. The onset of G1 initiates degradation of Ki67 (Miller et al., 2018). These cellular mechanisms have been considered to be part of the kidney’s adaptive repair process, which ultimately results in fibrosis (Nadasdy et al., 1994;Sheng and Zhuang, 2020).

In *Atxn10*^*Cagg*^ pancreata, the exocrine regions also exhibit similar actin misorganization, a loss of E-cadherin, up-regulation of Vimentin, and an increase in Ki67 positive cells. In the pancreas, the typical cellular response to an injury is through acinar-to-ductal metaplasia (ADM). Specifically, during ADM the epithelial pancreatic acinar cells are thought to assume a more progenitor or ductal epithelial cell status (Storz, 2017). Unlike the kidney, *Atxn10*^*Cagg*^ pancreata exhibit the added phenomena of an increase in ciliation in the exocrine region. In a normal pancreas, the cell types that are predominantly ciliated reside in the islet, and the ducts with the acini typically never possess a cilium (Augereau et al., 2016). Like *Atxn10*^*Cagg*^ kidneys, the pancreata also exhibit an increase in Ki67 positive cells. In both of these tissues, baseline proliferation is normally very low. It is possible that this prolonged proliferation stage in both the kidney and the pancreas is a failed repair process that would likely result in fibrosis if animals were to survive long enough.

The rapid decline in health of *Atxn10*^*Cagg*^ animals prevents the observation of longer term abnormalities. In the congenital model, the disposition towards cardiac abnormalities is likely due to the spatiotemporal expression pattern of *Atxn10*; however, in the inducible model it is currently unclear as to why specific tissues are preferentially effected. It is probable that if animals lived longer or if tissue specific Cre mouse lines were used, similar functions of ATXN10 in additional tissues would be uncovered. In the affected organs, the pancreatic acinar cells, renal epithelial tubules, and glandular epithelium are specifically disrupted. These epithelial cells begin to exhibit signs of disrupted polarity followed by loss of epithelial markers such as E-Cadherin and ZO-1. Concurrently, the mesenchymal marker, Vimentin, begins to be expressed in these cells indicating that they are likley undergoing EMT.

Collectively we show that ATXN10 is located at the base of the primary cilium, but it is not necessary for ciliogenesis. Furthermore, we show that ATXN10 is necessary for both embryonic and post-embryonic survival with loss in adult animals resulting in an EMT-like progression in the kidney and pancreas and with cells also undergoing ADM in the pancreas. In none of the adult induced mutants did we observe phenotypes consistent with SCA raising the possibility that the petanucleotide repeat in the SCA10 patients may not be due to loss of Atxn10 protein directly; however, the complication with this assessment is that the mice die rapidly following induction, and this may preclude the presentation of SCA phenotypes.

## Materials and Methods

### Generation of Atxn10 mutant alleles

All animal studies were conducted in compliance with the National Institutes of Health *Guide for the Care and Use of Laboratory Animals* and approved by the Institutional Animal Care and Use Committee at the University of Alabama at Birmingham. Mice were maintained on LabDiet^®^ JL Rat and Mouse/Irr 10F 5LG5 chow. The *Atxn10*^*KO*^ allele (*tm1a*) was rederived from sperm obtained from the Knockout Mouse Project (KOMP) Repository into C57/B6J strain mice. Mice were maintained on a mixed B6/129 background. *Atxn10* conditional allele (*tm1c*) mice were generated by mating the *Atxn10*^*KO*^ to FlpO recombinase mice (C57BL/6J) thus removing the LacZ and Neo cassettes and generating a conditional allele (tm1c; flox). Progeny that contained the recombined allele were crossed off of the FlpO line and bred to respective Cre recombinase males. Here we refer to these alleles as the *tm1a*(*Atxn10*^*KO*^), *tm1c*(*Atxn10*^*flox*^) and *tm1d*(*Atxn10*^*Cagg*^, *Atxn10*^*cTnT*^, or *Atxn10*^*Tie2*^) alleles. Primers used for genotyping are as follows: 5’-GACTTTTGGCACCACACAGC-3’, 5’-GTGGAAGGGCTGAAAACTGG-3’, 5’-TCGTGGTATCGTTATGCGCC-3’, 5’-ATCACGACGCGCTGTATC-3’, and 5’-ACATCGGGCAAATAATATCG-3’.

### Generation and transfection of ATXN10 expression constructs

The *MmAtxn10* coding sequence was cloned into the pEGFP-N1 vector (Clontech) using primers designed with XhoI and AgeI restriction sites. *MmAtxn10*::EGFP-N1 plasmids were transfected into cells using *Trans*IT^®^-2020 DNA per manufacturer guidelines (Mirus, MIR5404).

### Embryo Isolation

Timed pregnancies were established with embryonic time-point of E0.5 being noted at noon on the morning of observing the copulatory plug. To isolate embryos, pregnant females were anesthetized using isoflurane followed by cervical dislocation. Embryonic tissues or whole embryos were isolated and fixed in 4% paraformaldehyde (Sigma PFA, 158127) in PBS.

### β-Galactosidase staining

For whole mount or slice *β* -Galactosidase staining, samples were fixed (0.2% glutaraldehyde (Sigma), 5mM EGTA, 2mM MgCl_2_ in 1X PBS) at 4°C for 40 minutes. Samples were rinsed three times for 15 minutes at 4°C (0.02% Igepal, 0.01% Sodium Deoxycholate, and 2mM MgCl_2_ in 1X PBS). Samples were immersed in staining solution overnight in the dark at 37°C (1mg/ml X-gal, 0.02% Igepal, 0.01% Sodium Deoxycholate, 5mM potassium Ferricyanide, 5mM potassium Ferrocyanide, and 2mM MgCl_2_ in 1X PBS). Samples were post-fixed in 4% PFA and stored at 4°C. Embryos were imaged using a Nikon SMZ800 stereo microscope. Sections were counter stained using Nuclear Fast Red (Sigma).

### Isolation of Mouse Embryonic Fibroblasts (MEFs)

Embryos were isolated at either E9.5 (*Atxn10*^*KO/KO*^) or E13.5 (conditional lines). Following the removal of the liver (E13.5 only) and head, embryos were mechanically dissociated and cultured in DMEM (Gibco, 21063-021) supplemented with 10% Fetal Bovine Serum, 1X Penicillin and Streptomycin, 0.05% Primocin, 3.6μl/0.5L β-mercaptoethanol. Cilia formation was induced using media containing 0.5% FBS.

### Primary kidney epithelium cell culture

Mice were anesthetized with isofluorane followed by cervical dislocation. Kidneys were removed and mechanically disociated. Resulting minced tissue was filtered through a 70mm cell strainer. Tubules were cultured in DMEM (Gibco, 11039-021) supplemented with 5% FBS, Epidermal Growth Factor (recombinant human, 10ng/ml), Insulin (recombinant human, 5 μg/ml), Hydrocortisone (36ng/ml), Epinephrine (0.5 μg/ml), Triiodo-L-thyronine (4pg/ml), and Transferrin (recombinant human, 5 μg/ml) (Growth Medium 2 SupplementPack, PromoCell, C-39605).

### Pathology and histology

Mice were anesthetized with 0.1 ml/ 10 g of body weight dose of 2.0% tribromoethanol (Sigma Aldrich, St. Louis, MO) and transcardially perfused with PBS followed by 4% paraformaldehyde. Tissues were post-fixed in 4% PFA overnight at 4°C, cryoprotected by submersion in 30% sucrose in PBS for 16–24 hours, then embedded in OCT, and cryosectioned for immunofluorescence, and Hematoxylin (Fisher Chemical, SH26-500D) and Eosin (Sigma-Aldrich, HT110132-1L) staining was performed. Pathological and histological analyses for *Atxn10*^*Cagg*^ pancreata, kidneys, spleen, retina, lung, and liver were performed by the Comparative Pathology Lab (UAB) as follows. Briefly, mice were necropsied and tissues were fixed in 10% neutral buffered formalin overnight. Tissues were prosected and processed then 5 μM sections were stained with Hematoxylin and Eosin. Slides were evaluated for tissue histopathology by a board certified veterinary pathologist in blinded fashion.

### Immunofluorescence microscopy

Ten (10) μm tissue sections were used for immunofluorescence microscopy. For staining MEFs, cells were grown on 0.1% gelatin coated glass cover slips until confluent, then serum starved using DMEM containing 0.5% FBS for 24 hours to induce cilia formation (Breslow and Nachury, 2015). Sections were fixed with 4% PFA for 10 minutes, permeabilized with 0.1% Triton X-100 in PBS for 8 minutes and then blocked in a PBS solution containing 1% BSA, 0.3% TritonX-100, 2% (vol/vol) normal donkey serum and 0.02% sodium azide for one hour at room temperature. Primary antibody incubation was performed in blocking solution overnight at 4°C. Primary antibodies include: Acetylated α-tubulin (Sigma, T7451) direct conjugated to Alexa 647 (Invitrogen, A20186) and used at 1:1000, Arl13b (Proteintech, 1771-1AP, 1:500), PECAM1 (Abcam, ab7388, 1:250), E-Cadherin (Abcam,ab11512, 1:300), Phalloidin (Invitrogen, A12380 or A12379, 1:300) Ki67 conjugated to PE, (Thermofisher, 12-5698-80, 1:300) cTnT (DSHB, RV-C2, 1:300), ZO-1 (R40.76, 1:2), Vimentin (Abcam, ab92547, 1:300), Amylase (Abcam, ab189341, 1:1000) Sox9 (Abcam, ab185230, 1:300), Fluorescein labeled Lotus tetragobolobus lectin/LTA (Vector Laboratories, FL-1321, 1:250), Rhodamine labeled Dolichos Biflorus Agglutinin/DBA (Vector Laboratories, RL-1032, 1:250). Cryosections were washed with PBS three times for five minutes at room temperature. Secondary antibodies diluted in blocking solution were added for one hour at room temperature. Secondary antibodies included: Donkey conjugated Alexa Fluor 647, 488, and 594 (Invitrogen, 1:1000). Samples were washed in PBS and stained with Hoechst nuclear stain 33258 (Sigma-Aldrich) for 5 minutes at room temperature. Cover slips were mounted using SlowFade Diamond Antifade Mountant (Life Technologies). All other fluorescence images were captured on Nikon Spinning-disk confocal microscope with Yokogawa X1 disk, using Hamamatsu flash4 sCMOS camera. 60x apo-TIRF (NA=1.49), 40x plan flour (NA=1.3), or 20x Plan Fluor Multi-immersion (NA=0.8) objectives were used. Images were processed using Nikon’s Elements or Fiji software.

### Tamoxifen Cre induction

Recombination of the conditional allele was induced in *Atxn10*^*flox/flox*^;*CAGG-cre*^*ERT2*^ mice at 6 and 8 weeks old by a single intraperitoneal (IP) injection of 9mg tamoxifen (Millipore Sigma, T5648) per 40g (body weight) in corn oil. Induction of cell lines was achieved by exposure to media supplemented with 1mM 4-hydroxytamoxifen for 24h.

### Statistics

Calculations were performed using Graphpad Prism and Microsoft Excel. Specific tests used are indicated in figure legends with significance indicated as follows: * p≤0.05, ** p≤0.01, *** p≤0.001. Error bars indicate Standard Error of the Mean (SEM).

## Acknowledgments

The authors would like to thank the members of Dr. Bradley K. Yoder’s (UAB) and Dr. Jeremy F. Reiter’s (UCSF) laboratories for intellectual and technical support on the project. The authors would like to thank the UAB Comparative Pathology Lab and Jeremy Foote, DVM, PhD for his expertise. The authors would like to thank the National Institute of Child Health and Human Development and the National Heart, Lung, and Blood Institute for financial support of these studies.

---

**Supplemental Figure 1.**
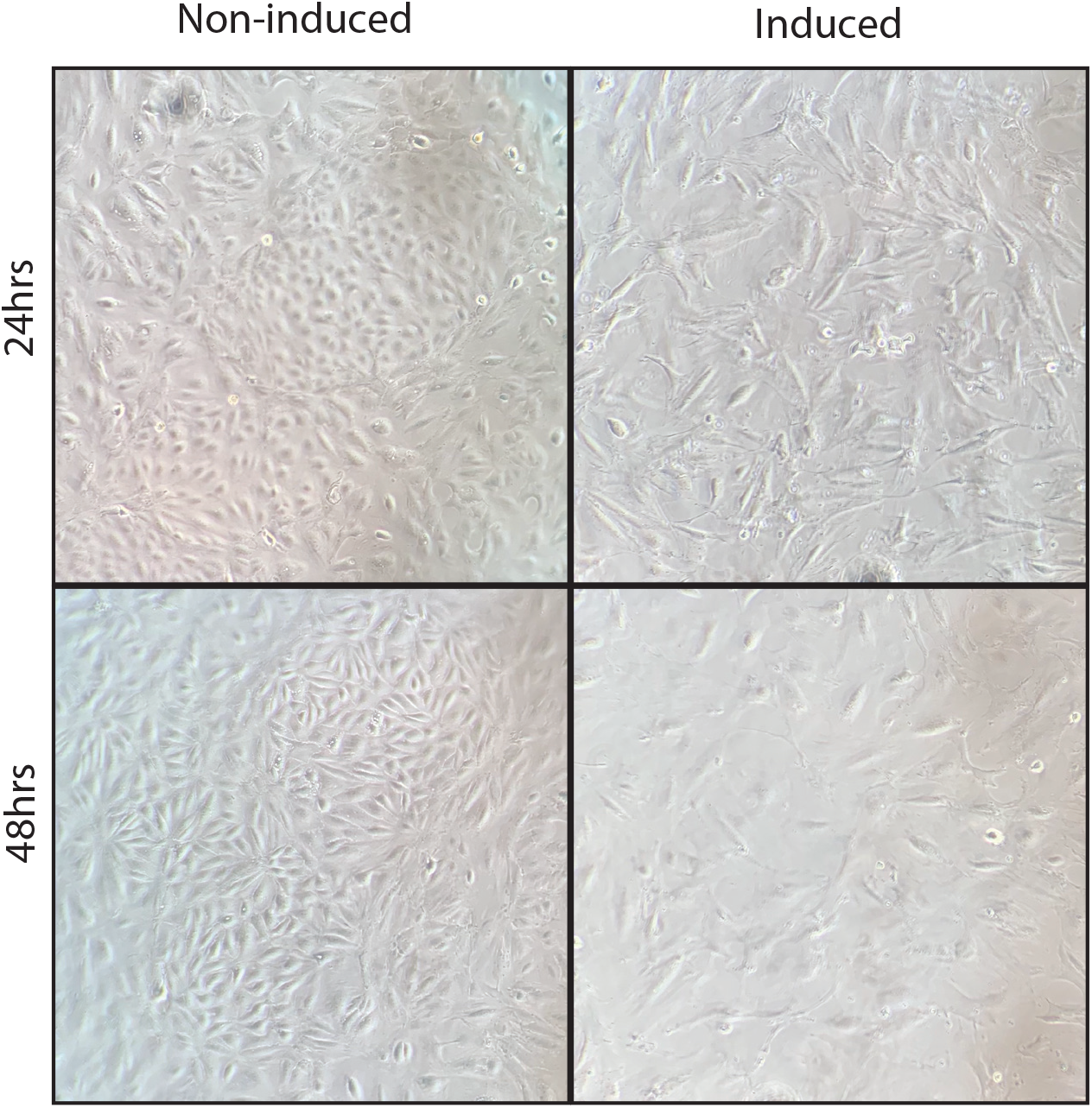
Primary kidney epithelial cells post induction. Phase contrast image of induced (right) and non-induced (left) cultured primary kidney epithelial cells from Atxn10^flox/flox^; Cagg-Cre^ERT^ mouse kidneys. Top panels indicate cells 24 hours post induction and bottom panels indicate cells 48 hours post induction.

**Supplemental Figure 2.**
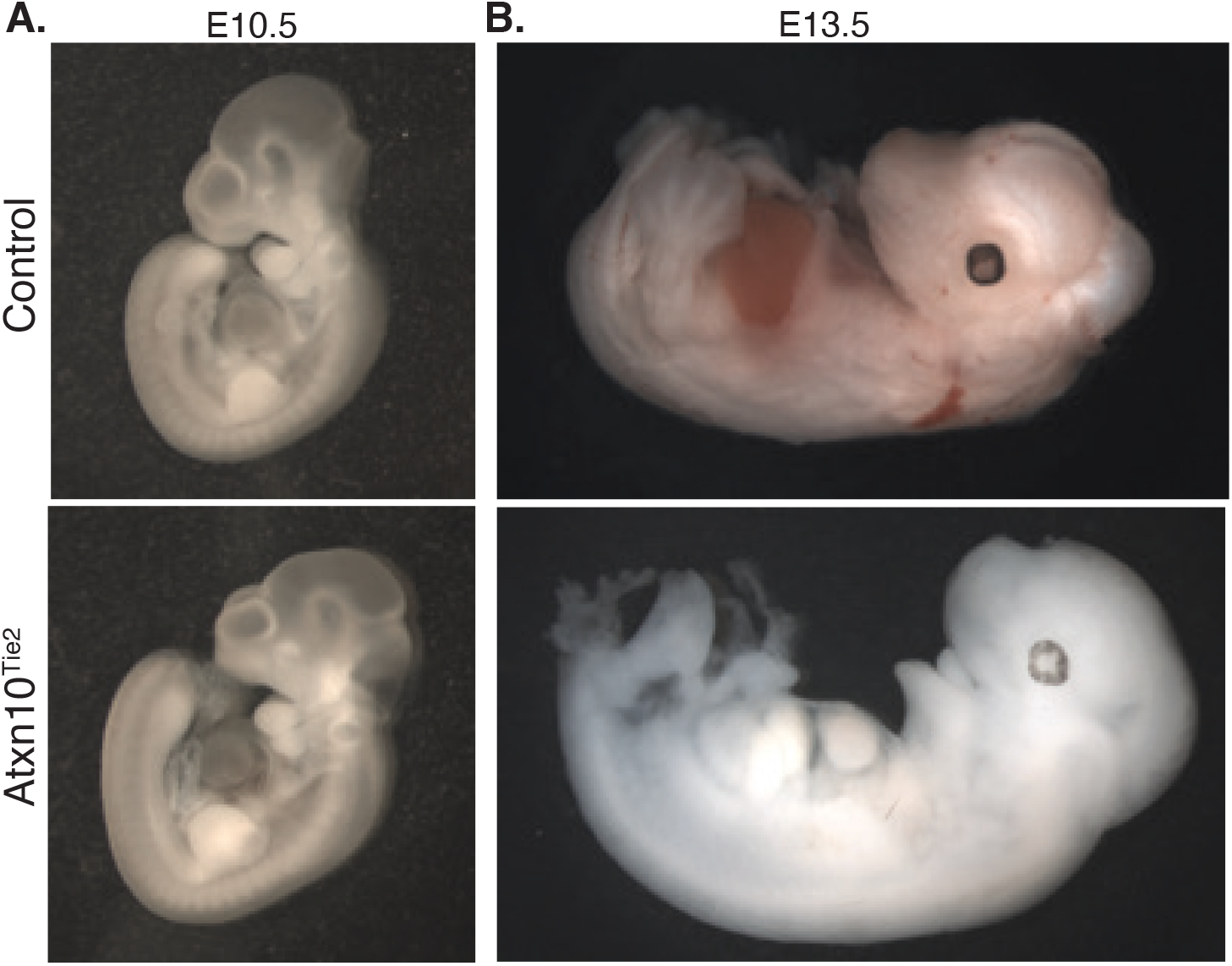
*Atxn10*^*Tie2*^ embryos. Control (top) and *Atxn10*^*Tie2*^ (bottom) embryos at A) E10.5 (left) and B) E13.5 (right).

**Supplemental Figure 3.**
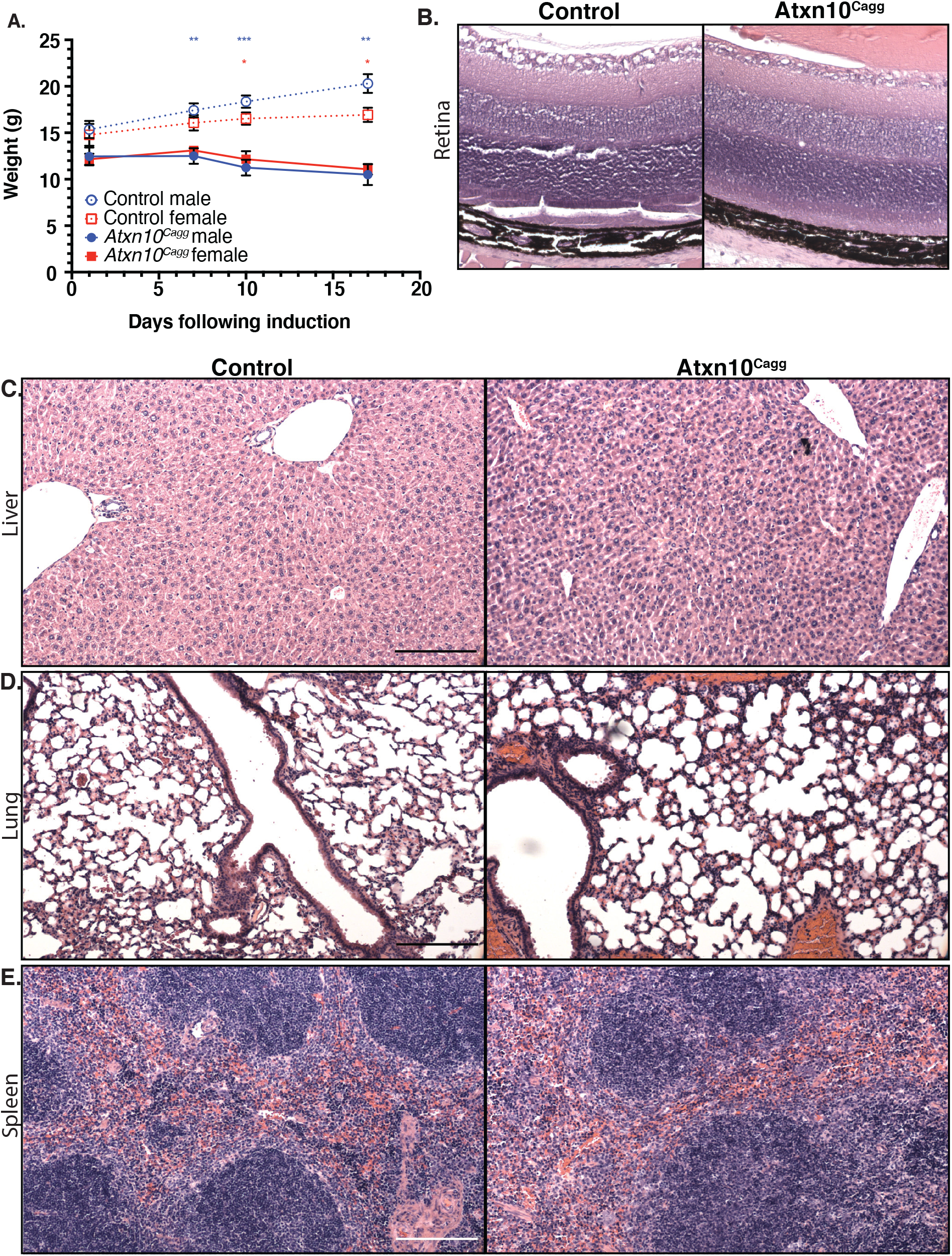
*Atxn10*^*Cagg*^ unaffected tissues. A) Weights following induction of *Atxn10*^*flox/flox*^ and *Atxn*10^Cagg^ animals at 4 weeks old (Statistical significance is designated by red asteriks for females and blue asteriks for males; N= 5 Cre- females, 6 Cre- males, 4 Cre+ females, and 6 Cre+ males). H&E staining of Control (left) and *Atxn*10^Cagg^ (right) B) retina, C) liver, which demonstrates hepatocyte atrophy consistent with anorexia and weight loss D) lung, E) spleen. Scale bars=50μm. Statistical significance was determined using mixed effects analysis with multiple comparisons.

**Supplemental Figure 4.**
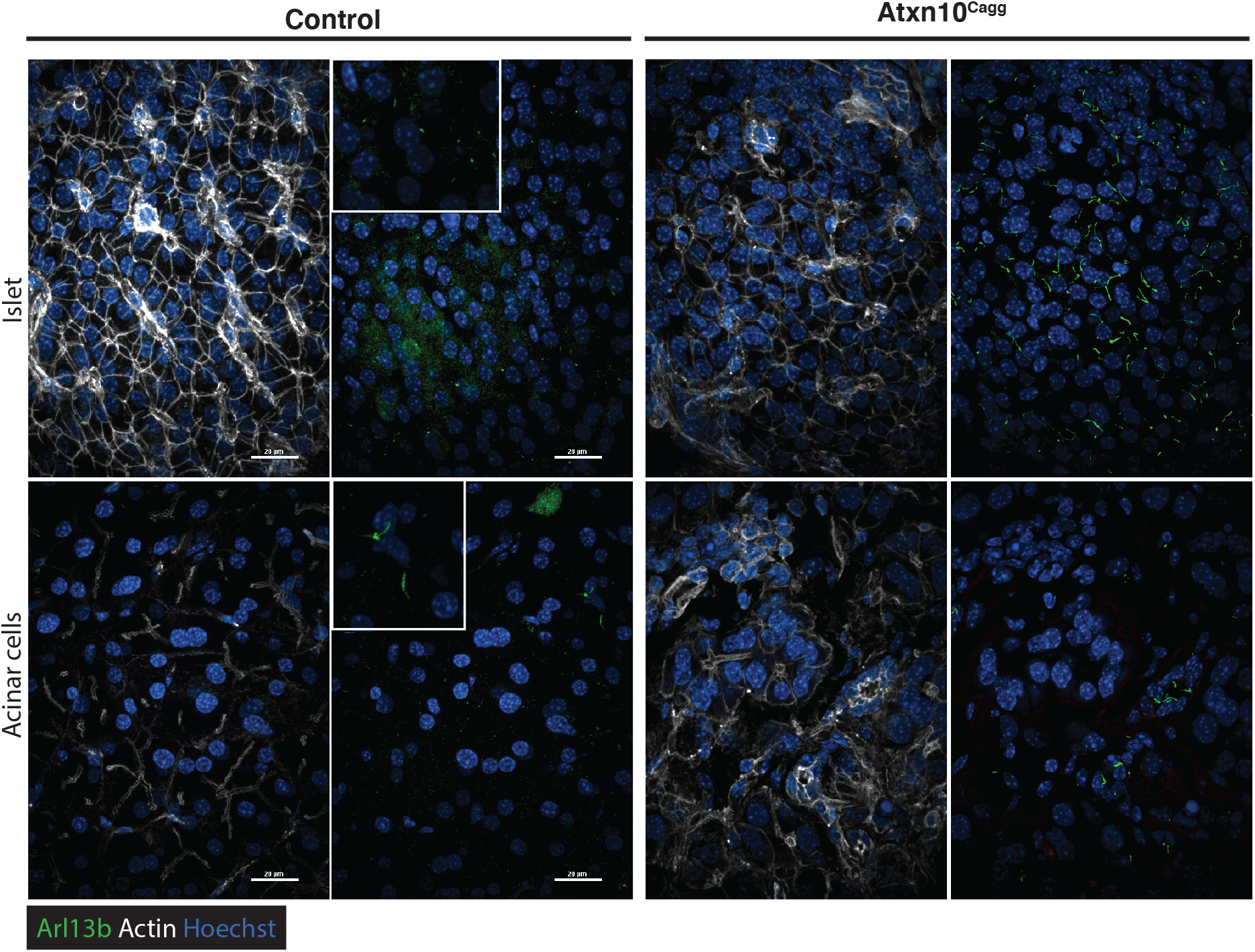
Ectopic cilia in *Atxn10*^*Cagg*^ pancreas. Immunofluorescence staining for Actin (white), cilia shown by Arl13b staining (green), and Hoechst (blue) in Control and *Atxn*10^Cagg^ islet (top) and acinar cells (bottom) in the pancreas (scale bar= 20 μm).

